## Supplementary material for "Variation in the ACE2 receptor has limited utility for SARS-CoV-2 host prediction"

#### Supplementary files

**Supplementary file 1.** Final data on susceptibility to, and shedding of, sarbecoviruses, along with the accession numbers for ACE2 amino acid sequences used to represent each species. These data were used to train all models presented.

**Supplementary file 2.** Predictions from the phylogeny-only model. Predictions are separated into four categories, reflecting predictions for (a) species in the training data, produced by withholding each species in turn from model training (i.e., predictions informing the leave-one-out cross-validation statistics presented), (b) susceptible species recognised through natural infection after data collection for this study had ended, (c) all other mammals available in the TimeTree phylogeny, and (d) all other birds in the TimeTree phylogeny. Across all tables, “cutoff” refers to the optimised value beyond which predicted probabilities from a given model are considered to indicate susceptible species (labelled as “True” in the “prediction” column).

### Figure supplements

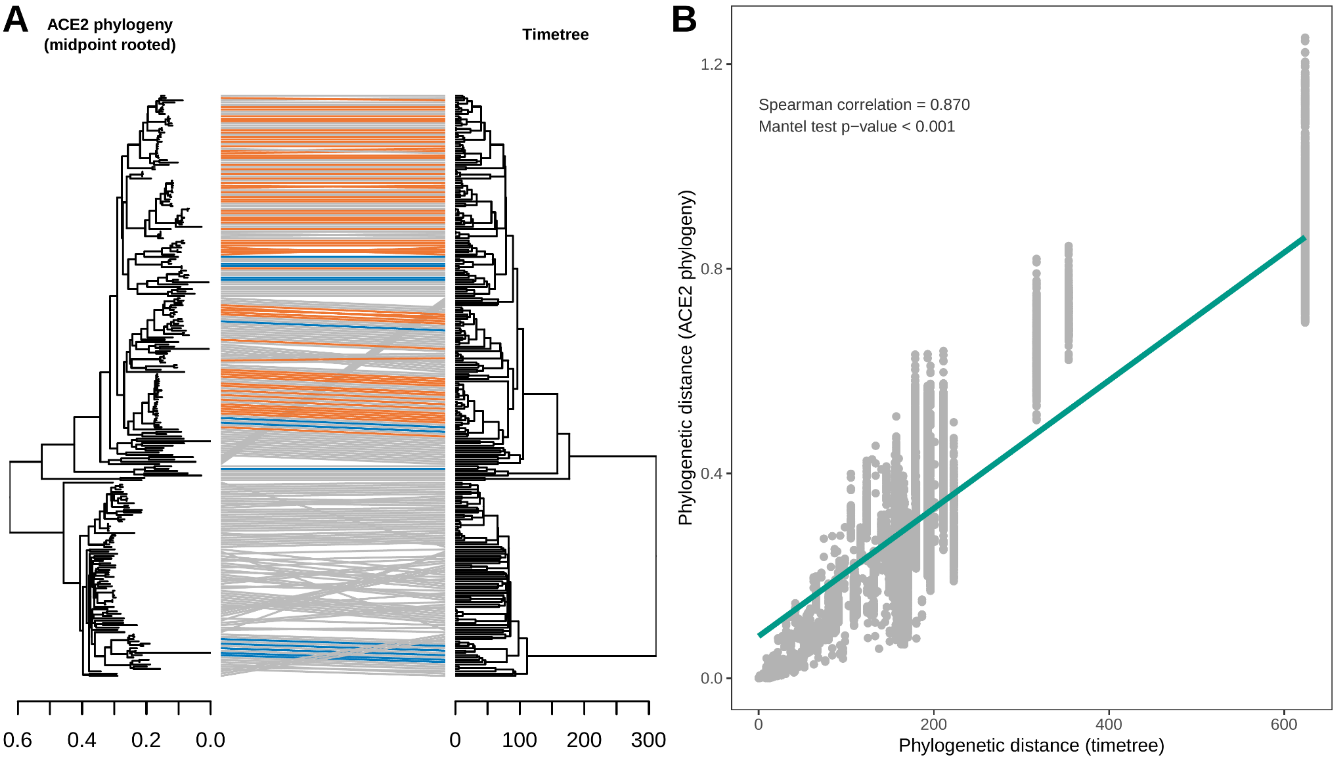


**Figure 1 – figure supplement 1.** Congruence between a phylogeny reconstructed from ACE2 amino acid sequences and a consensus time-scaled phylogeny for mammals and birds obtained from the TimeTree database. (A) Tanglegram linking species across the two phylogenies, with sarbecovirus-susceptible species indicated in orange and putatively non-susceptible species in blue. Grey lines indicate species with unknown susceptibility. The maximum likelihood ACE2 phylogeny was constructed using an alignment of all available ACE2 ortholog sequences from species also occurring in the TimeTree database (N=253). The TimeTree phylogeny was trimmed to contain the same species. (B) Correlation between pairwise cophenetic distances calculated from the two phylogenies. A linear regression trendline is shown for visualisation purposes only.

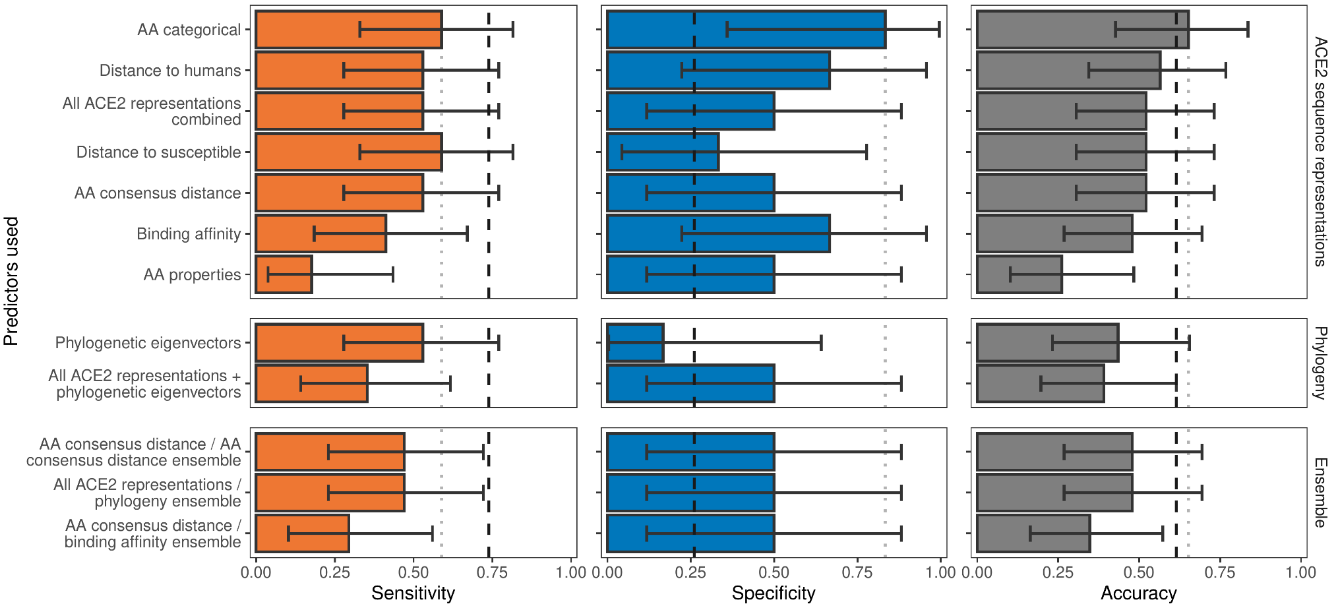


**Figure 2 – figure supplement 1**. Ability of models trained on different representations of either ACE2 sequences or a time-scaled amniote phylogeny to predict shedding of infectious virus after sarbecovirus infection. Bars represent proportions from leave-one-out cross-validation, with error bars indicating binomial confidence intervals. Dashed vertical lines indicate the performance expected from a null model which randomly assigns shedding / non-shedding labels in proportion to their frequency in the training data (73.9% of hosts were reported to shed infectious virus, N = 23). Dotted vertical lines highlight performance of the best model in each panel. The low number of species for which their ability to shed infectious virus has been reported means no tested model currently outperforms random guessing across both sensitivity and specificity.

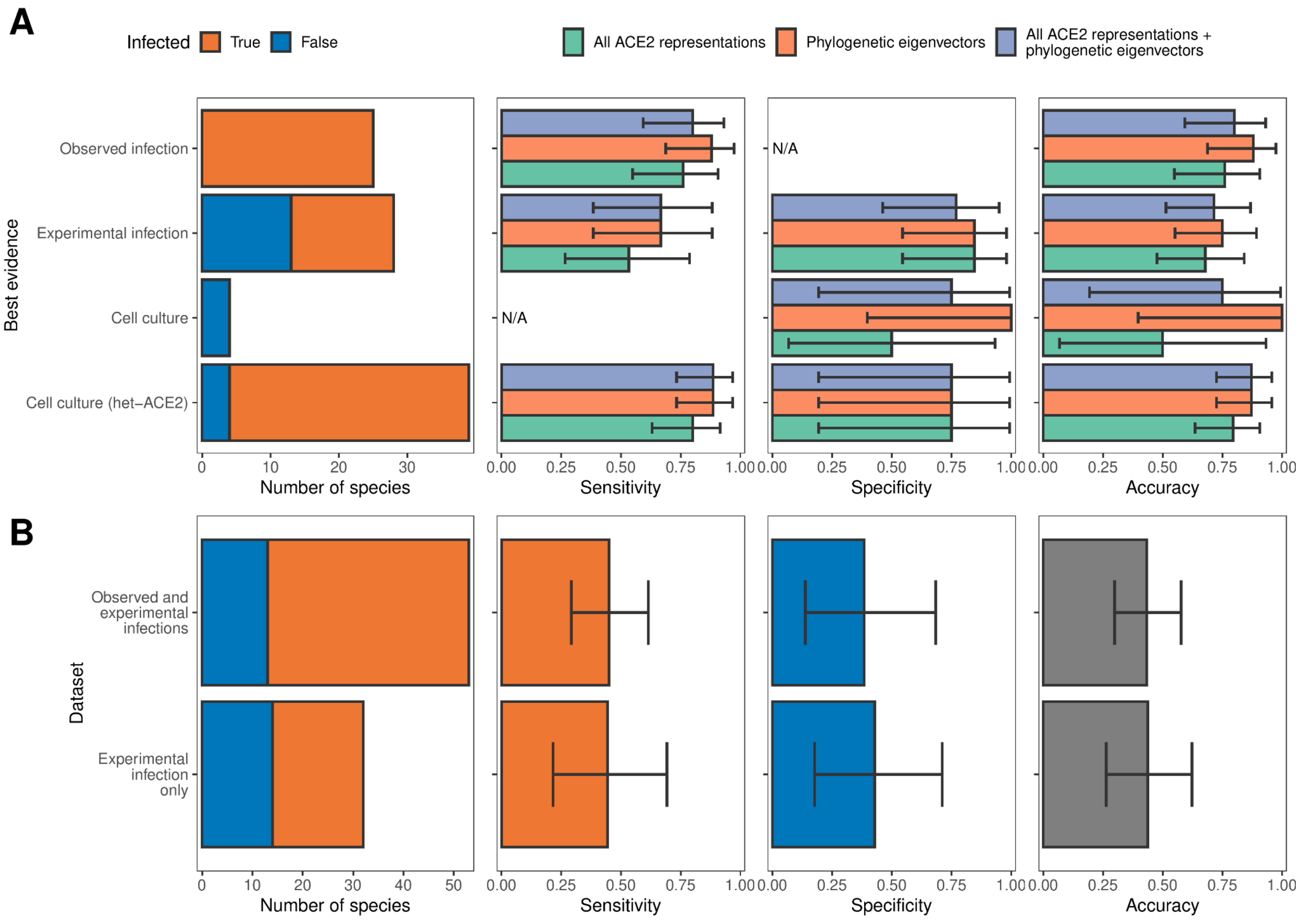


**Figure 2 – figure supplement 2.** Influence of different sources of sarbecovirus susceptibility data on prediction accuracy. (A) Performance of different models trained on all available data. Holdout predictions for individual species were grouped by the best available evidence of susceptibility or non-susceptibility to calculate performance metrics only. N/A indicates no suitable observations in a given category (e.g., cell culture data added no new susceptible species, meaning we cannot calculate the sensitivity of models on cell culture data). These are the same models as in Figure 2. (B) Performance of two alternative models trained using all ACE2 representations combined with phylogenetic eigenvectors, but with subsets of the full dataset as training data. The first panel in both rows indicates the size of each data subset. In all other panels, bars represent proportions from leave-one-out cross-validation, with error bars indicating binomial confidence intervals. Since there are no consistent trends in model performance across data sources in (A), we conclude that the inclusion of cell culture and heterologous ACE2-based data are not affecting our ability to compare different models. In contrast, removing these data sources leads to severe decreases in model performance, likely due to low sample sizes (B). This identifies cell culture experiments, including experiments testing only binding of heterologous ACE2, as useful additional sources of compatibility data when this would otherwise be limited.

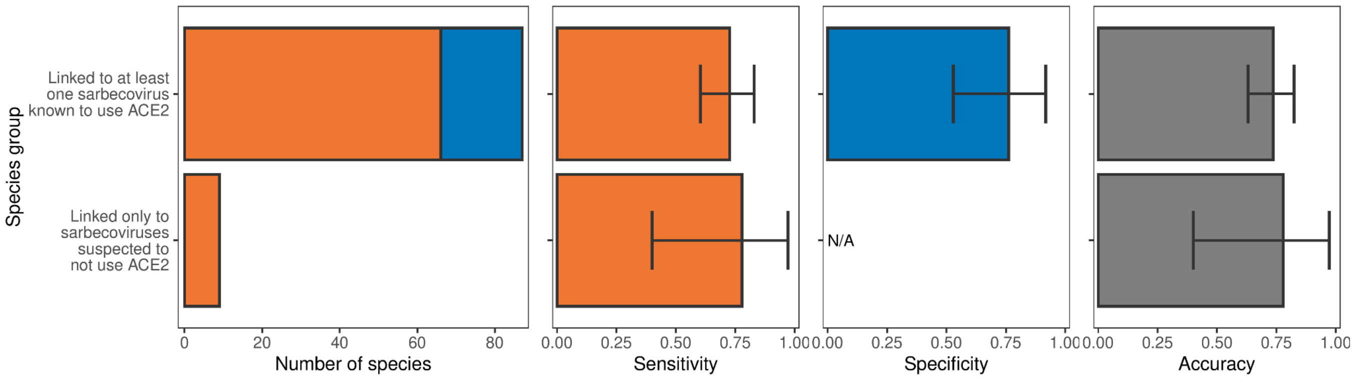


**Figure 2 – figure supplement 3**. Performance of a model trained with all ACE2-representations on hosts linked to sarbecoviruses not known to use ACE2. The model was trained on all hosts, with performance evaluated using leave-one-out cross-validation. In the first panel, bars represent sample sizes for susceptible species (orange) and non-susceptible species (blue). In all other panels, bars represent the proportion of species in each category accurately predicted, with error bars indicating binomial confidence intervals. Similar performance suggests ACE2-based models are equally capable of predicting the susceptibility of species thus far linked only to non-ACE2-utilising sarbecoviruses and those linked to other sarbecoviruses.

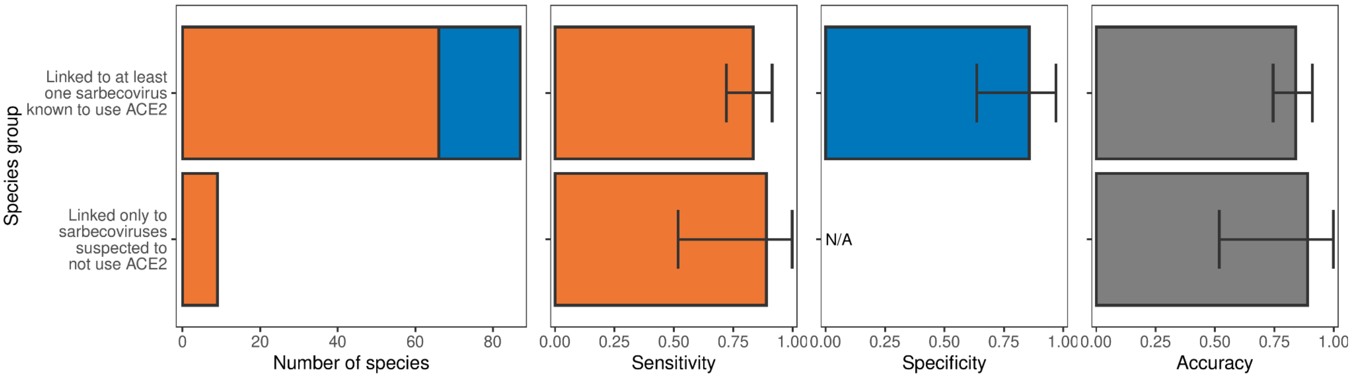


**Figure 2 – figure supplement 4.** Performance of a model trained with phylogenetic eigenvectors on hosts linked to sarbecoviruses not known to use ACE2. The model was trained on all hosts, with performance evaluated using leave-one-out cross-validation. In the first panel, bars represent sample sizes for susceptible species (orange) and non-susceptible species (blue). In all other panels, bars represent the proportion of species in each category accurately predicted, with error bars indicating binomial confidence intervals. As was the case for ACE2-based models (Figure 2 – figure supplement 3), phylogeny-based models are equally capable of predicting the susceptibility of species thus far linked only to non-ACE2-utilising sarbecoviruses and those linked to other sarbecoviruses.

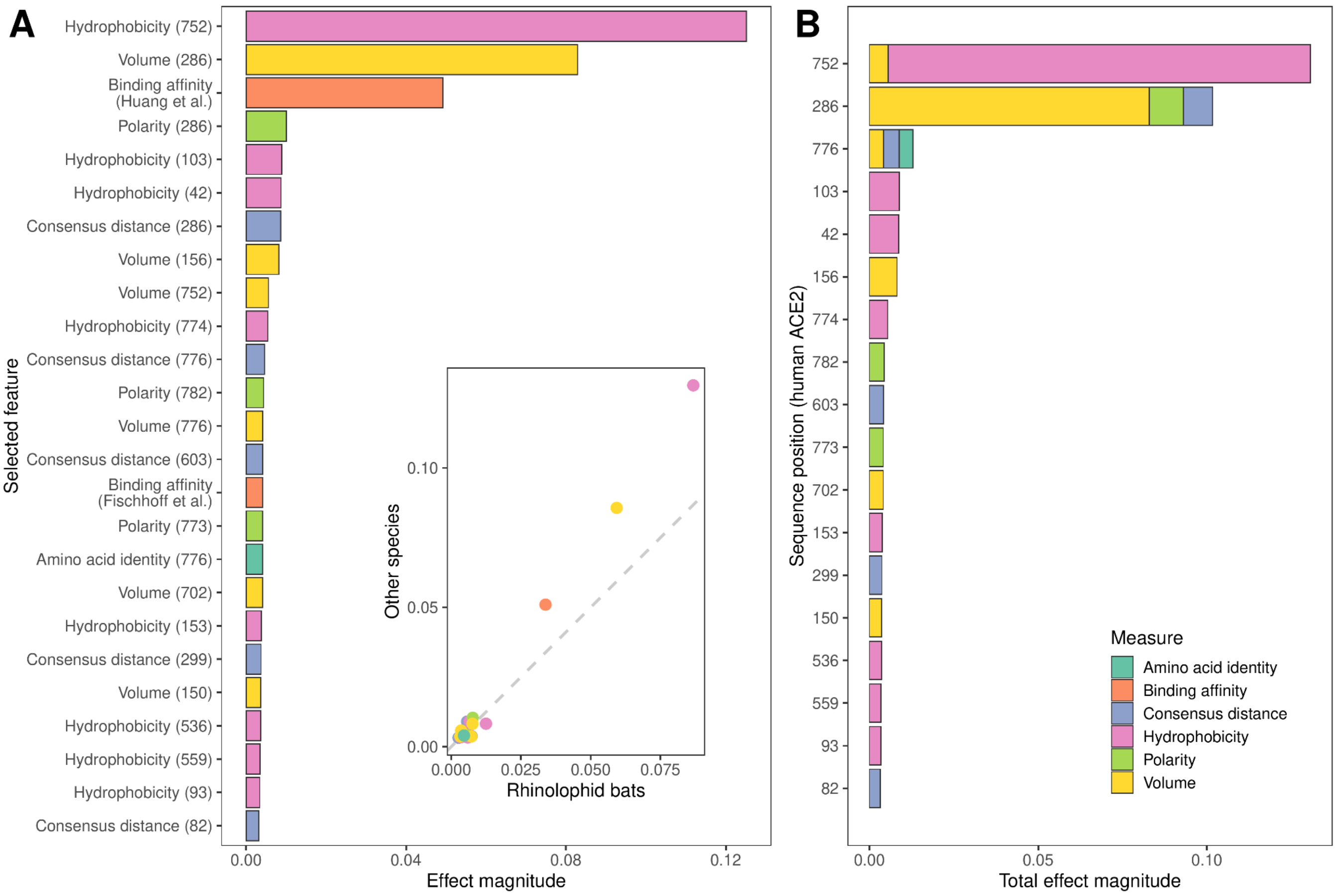


**Figure 3 – figure supplement 1.** Features retained and used by the combined ACE2-based model for predicting susceptibility to sarbecovirus infection. (A) Importance of individual features in determining final predictions. Importance was measured using mean absolute SHAP values across all host species in the training data. An inset shows the same mean effect magnitudes calculated separately for rhinolophid bats and other species, with a dashed line indicating a 1:1 relationship. Both axes of the inset plot are shown on a $log_{10}$ scale for clarity. Since all effect magnitudes are similar between rhinolophid bats and other species (indicated by being close to the dashed line), we conclude that the ACE2-based model trained on all species uses the same limited number of features and sequence positions to predict susceptibility of both groups. (B) Combined importance for individual ACE2 sequence positions, obtained by summing values from (A) whenever features represent the same position. Sequence positions refer to locations in the human ACE2 reference sequence (accession number NP_001358344.1).

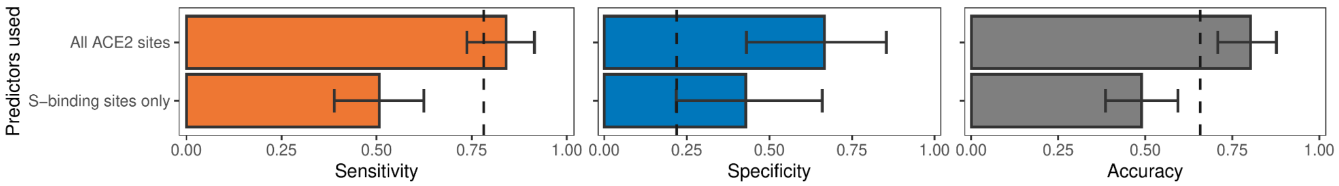


**Figure 3 – figure supplement 2.** Model performance when training the best-performing non-ensemble ACE2-only model ("AA consensus distance") with access to either all sites (as in Figure 2), or with representations of known SARS-CoV-2 spike-binding sites only. Both models represent individual ACE2 sites as a distance between the observed amino acid and the most common amino acid at that site among known susceptible species. Spike-binding sites were identified from annotations to the human ACE2 sequence (accession number NP_001358344.1). Removing phylogenetically-informative sites to focus on ACE2 positions with known mechanistic links to susceptibility decreases our ability to predict susceptibility to sarbecovirus infection.

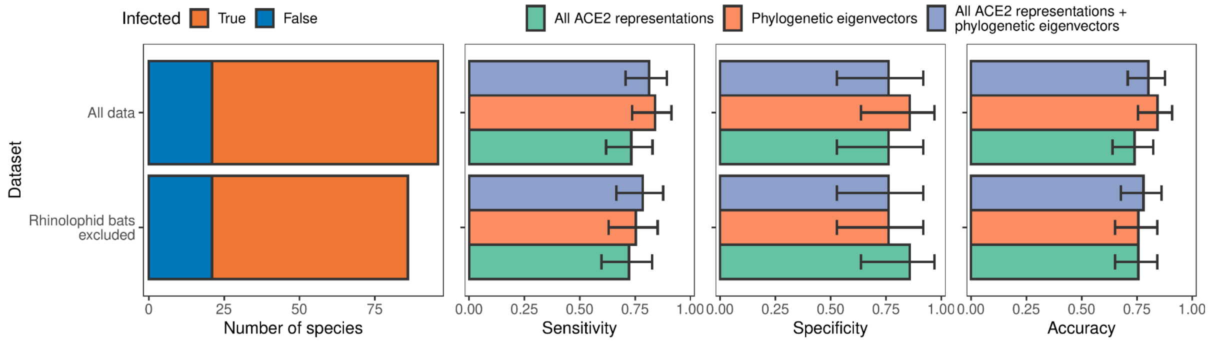


**Figure 3 – figure supplement 3.** Model performance after training with and without data from rhinolophid bats. Excluding rhinolophid bats has no effect on model performance, suggesting that any potentially-different evolutionary pressures on the ACE2 sequences of putative sarbecovirus reservoirs compared to other species did not negatively affect our models.

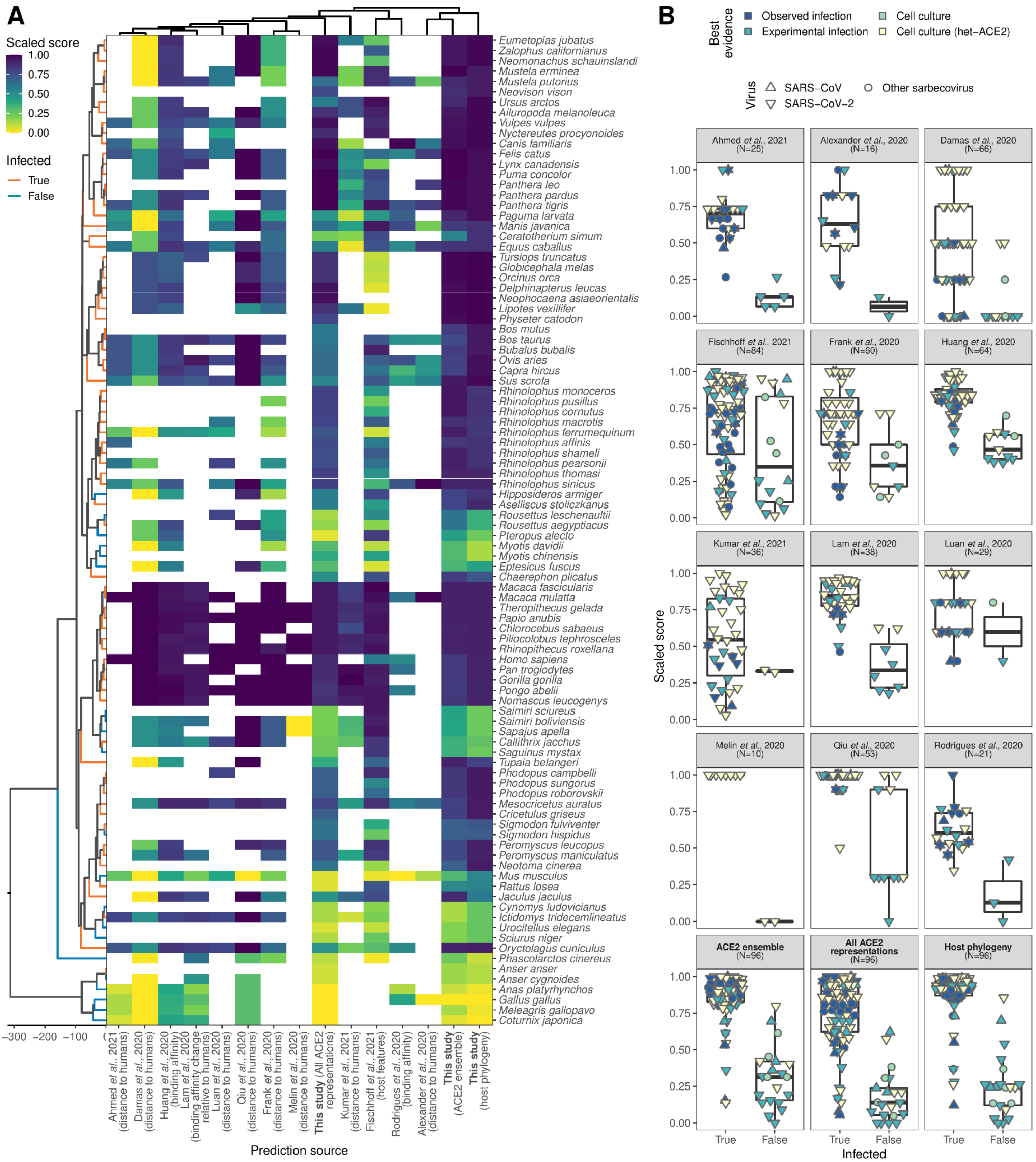


**Figure 4 – figure supplement 1.** Comparison of predictions across studies and models. (A) Heatmap comparing overlapping quantitative predictions for the same species across studies, where susceptibility is known. Quantitative scores from all studies were re-scaled to lie in [0, 1] to allow comparison. A time-scaled phylogeny along the left edge shows evolutionary relationships between species (derived from TimeTree, scale bar indicates millions of years), while the hierarchical clustering dendrogram above the heatmap shows similarities between studies based on pairwise Spearman correlations (ρ) across all overlapping species (including species with unknown susceptibility). (B) Ability of different heuristics to separate known susceptible and non-susceptible species. Boxplots were calculated using one observation per species, but overlapping symbols at the same position are used to indicate known susceptibility or non-susceptibility to multiple viruses. To allow comparison with published heuristics, the displayed quantitative scores from this study were taken from models fitted to all data, meaning they represent fitted values rather than holdout predictions. This matches the way quantitative scores were produced in earlier heuristics; all other plots use holdout predictions. Clustering of our “All ACE2 representations” model in the midst of previously-published heuristics (panel A, maximum $\rho = 0.744$) confirms that our ACE2-based models capture the bulk of currently-used approaches for predicting sarbecovirus host range as intended, meaning our formal holdout-based results showing an inability of ACE2-based models to outperform phylogeny likely extends to these earlier heuristics (also visible in (B)).

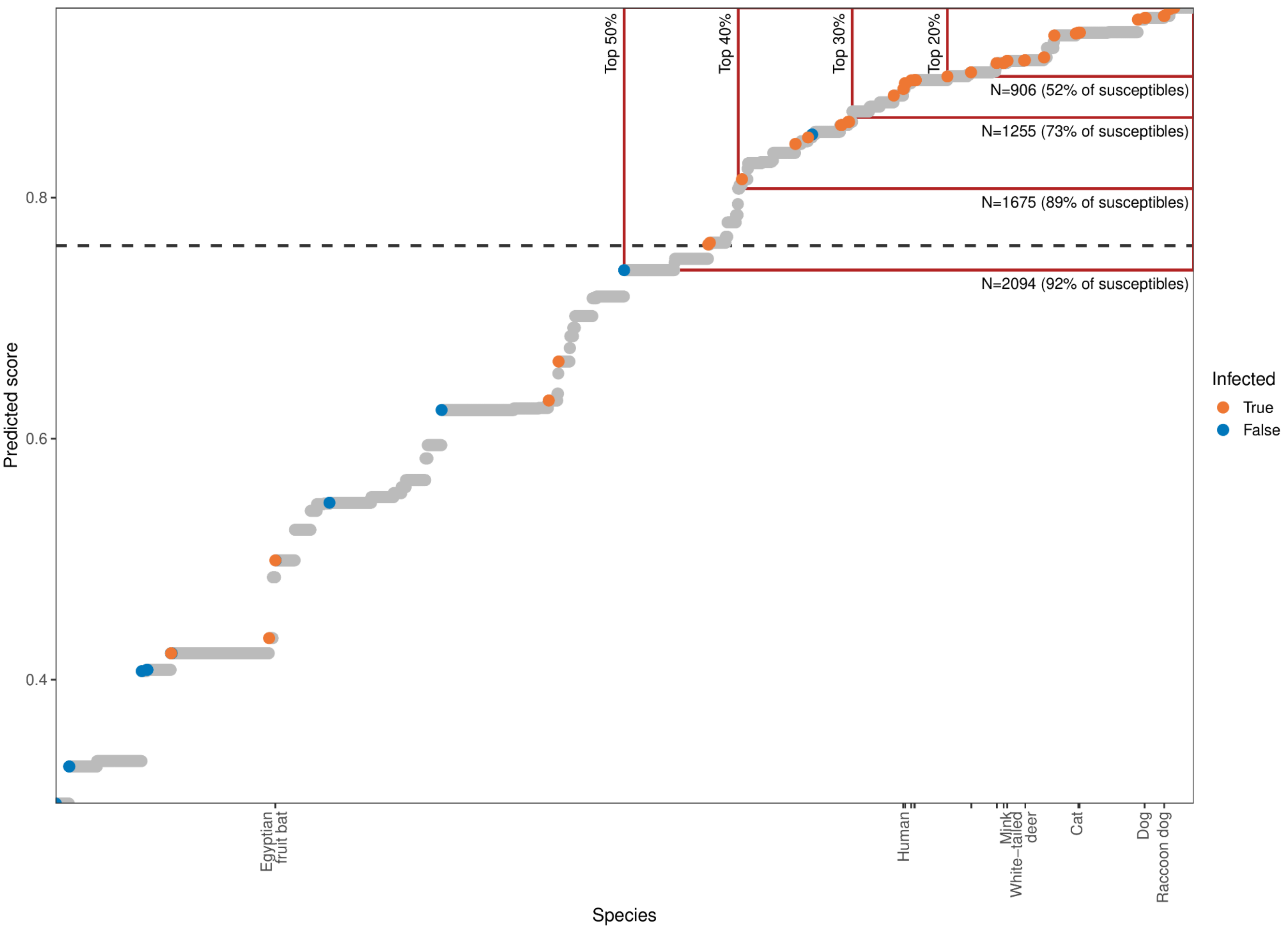


**Figure 5 – figure supplement 1.** Estimating the value of quantitative susceptibility predictions from the phylogeny-only model for guiding surveillance. Mammal species are arranged by model output, with higher scores indicating species predicted as more likely to be susceptible to sarbecovirus infection. A dashed line shows the optimised cut-off beyond which species were predicted as susceptible. Colours indicate known susceptible or non-susceptible species. Tick marks along the x-axis indicate the location of species known to also shed infectious virus after infection, with a limited selection of species labelled for orientation. Red boxes highlight the effect of focusing surveillance effort on the top-ranked species only, with the percentage of all species included indicated by vertical labels. Horizontal labels highlight the number of mammal species included by each cut-off, with the proportion of all known susceptible species included indicated in parentheses. Attempts to limit the number of species requiring surveillance to feasible numbers by focussing only on the most confident predictions risks missing large numbers of susceptible species.

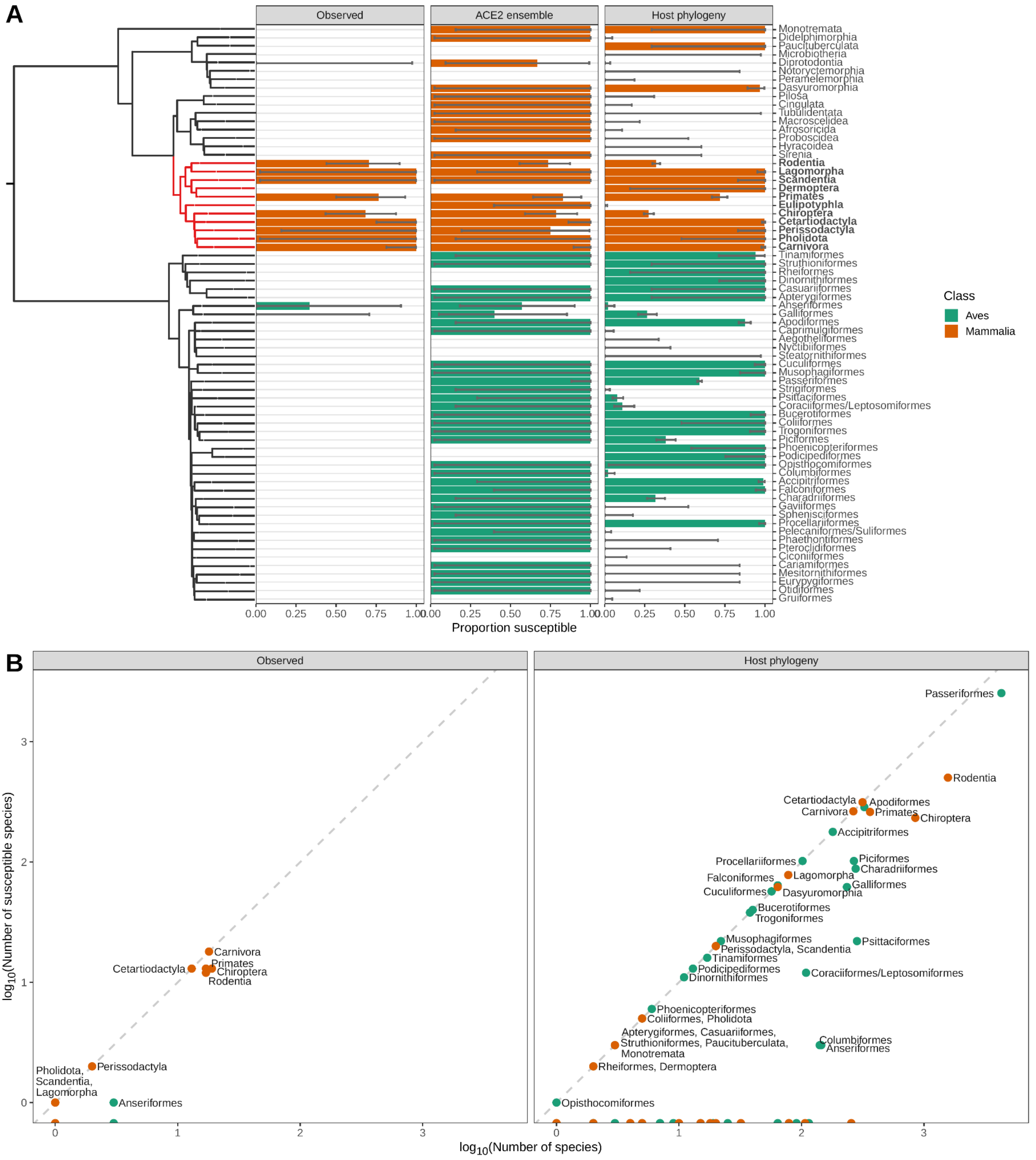


**Figure 5 – figure supplement 2.** Species observed and predicted as susceptible to sarbecovirus infection, aggregated by taxonomic order. (A) Proportion of species either observed or predicted to be susceptible by different models. Error bars indicate binomial confidence intervals. A time-scaled phylogeny illustrates divergence patterns between groups (derived from TimeTree). Taxonomic orders which were polyphyletic in this phylogeny were merged, and are indicated using a forward-slash (e.g., Coraciiformes/Leptosomiformes). Red branches and bold labels highlight the Boreoeutheria clade. (B) Relationship between the number of susceptible species in each taxonomic order and the size of each order. Models can exclude some taxonomic orders from requiring surveillance, but most orders which do contain susceptible species show limited variation in susceptibility. This limits our ability to target surveillance below the taxonomic order level (but see Figure 5 – figure supplement 3).

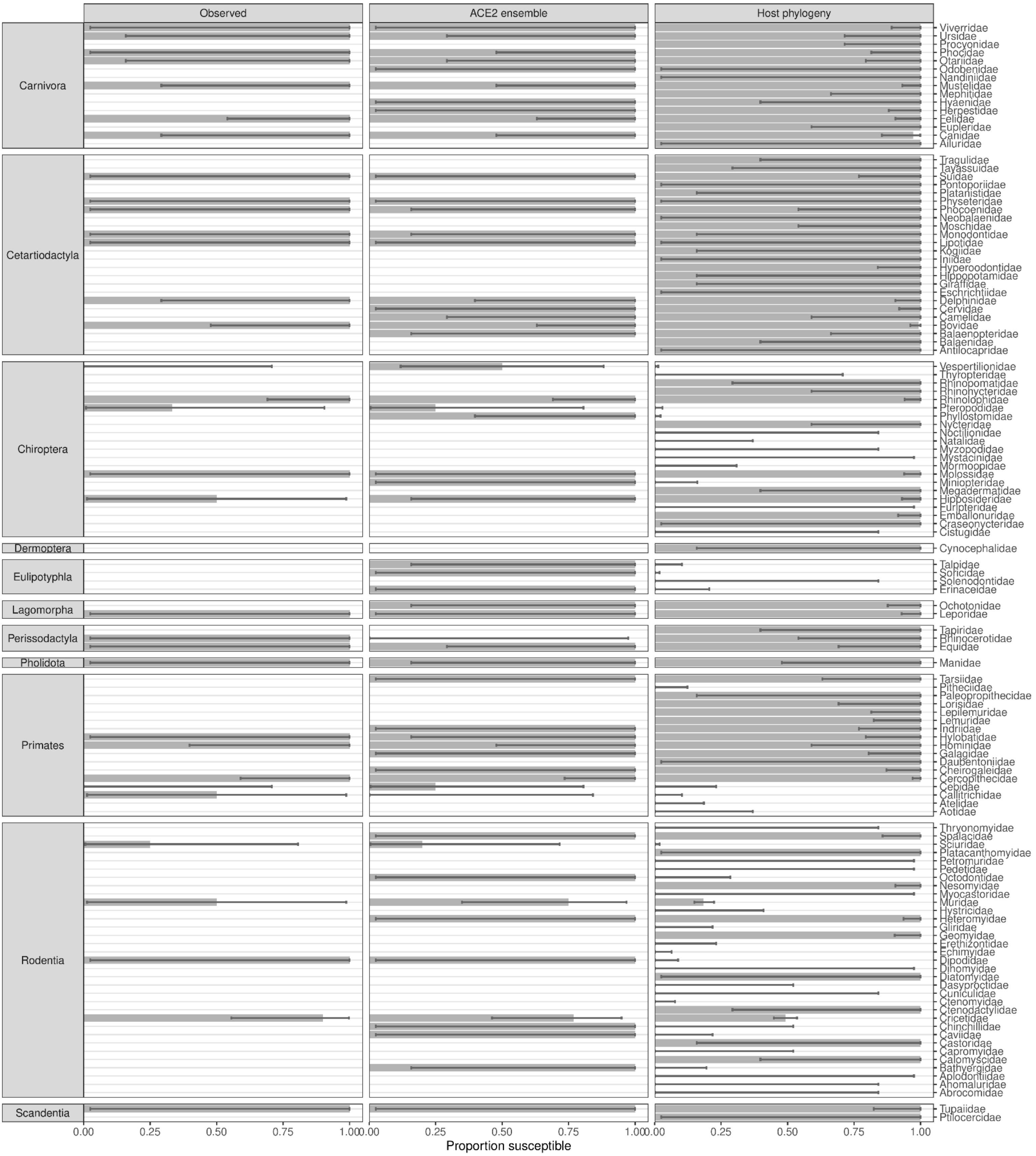


**Figure 5 – figure supplement 3.** Proportion of species observed or predicted to be susceptible to sarbecovirus infection in boreoeutherian families. All taxonomic orders within the Boreoeutheria clade containing at least one observed or predicted susceptible species are shown. Error bars indicate binomial confidence intervals. The Chiroptera and Rodentia orders include several families predicted to contain no susceptible species, but both current observations and model predictions suggest little variation in susceptibility below the family level.

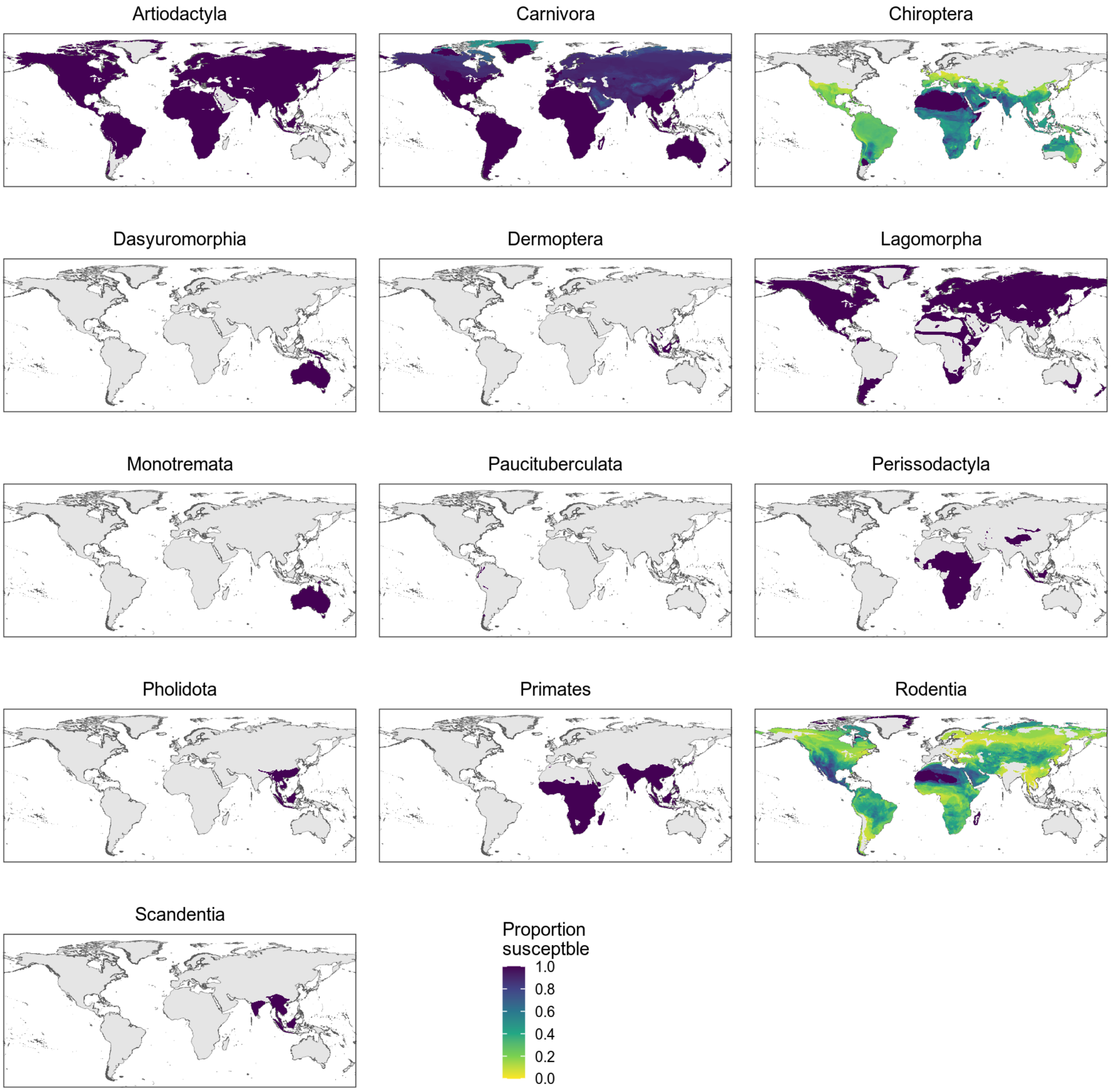


**Figure 5 – figure supplement 4**. Distribution of wild terrestrial mammals predicted as susceptible by the phylogeny-only model, separated by taxonomic order. All taxonomic orders within the Boreoeutheria clade containing at least one species predicted as susceptible are shown. Only bats and rodents show variation, likely due family-level differences in susceptibility (Figure 5 – figure supplement 3).
